# The Role of ATP Hydrolysis and Product Release in the Translocation Mechanism of SARS-CoV-2 NSP13

**DOI:** 10.1101/2023.09.28.560057

**Authors:** Monsurat M. Lawal, Priti Roy, Martin McCullagh

## Abstract

In response to the emergence of COVID-19, caused by SARS-CoV-2, there has been a growing interest in understanding the functional mechanisms of the viral proteins to aid in the development of new therapeutics. Non-structural protein 13 (Nsp13) helicase is an attractive target for antivirals because it is essential for viral replication and has a low mutation rate; yet, the structural mechanisms by which this enzyme binds and hydrolyzes ATP to cause unidirectional RNA translocation remain elusive. Using Gaussian accelerated molecular dynamics (GaMD), we generated a comprehensive conformational ensemble of all substrate states along the ATP-dependent cycle. ShapeGMM clustering of the protein yields four protein conformations that describe an opening and closing of both the ATP pocket and RNA cleft. This opening and closing is achieved through a combination of conformational selection and induction along the ATP cycle. Furthermore, three protein-RNA conformations are observed that implicate motifs Ia, IV, and V as playing a pivotal role in an ATP-dependent inchworm translocation mechanism. Finally, based on a linear discriminant analysis of protein conformations, we identify L405 as a pivotal residue for the opening and closing mechanism and propose a L405D mutation as a way of testing our proposed mechanism. This research enhances our understanding of nsp13’s role in viral replication and could contribute to the development of antiviral strategies.

**TOC Graphic:** 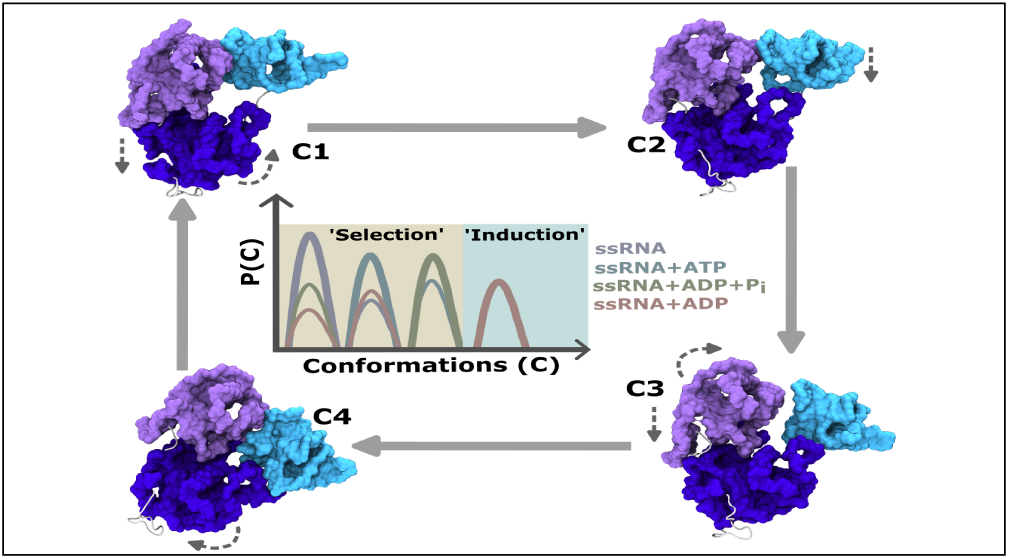

## Introduction

The severe acute respiratory syndrome coronavirus 2 (SARS-CoV-2) is the causative agent of the COVID-19 pandemic, which remains a global public health concern.^1^ Currently, COVID-19 has claimed the lives of nearly 7 million people worldwide, with over 1 million deaths reported in the USA as of the manuscript’s writing. ^2^ While vaccination has proven effective in mitigating severe disease,^3^ SARS-CoV-2 a positive-sense single-stranded RNA (+ssRNA) virus continues to undergo genetic changes, leading to the emergence of new variants that require evolving efforts for mitigation. ^4,5^ The SARS-CoV-2 virus possesses a genome encoding sixteen non-structural proteins (NSPs), seven accessory proteins, and four structural proteins.^6^ Nsp13 is a pivotal, yet under studied, NSP that acts primarily as a helicase during viral replication,^7^ but is also involved in RNA 5*′*-triphosphatase activity^8,9^ and mRNA capping.^10^ During viral RNA replication, at least one nsp13 interacts with the viral RNA-dependent RNA polymerase (RdRp) nsp12^11^ and other NSPs (nsp7 and nsp8) to form the replication-transcription complex.^11–14^ With the emergence of new variants, this protein displays a low mutation rate,^15^ underscoring its critical role in the virus’s life cycle. In this study, we present fundamental research into the functional mechanisms of nsp13, a SARS-CoV-2 protein, which can be harnessed for the development of new antiviral treatments.

SARS-CoV-2 nsp13 is classified as a member of the superfamily 1 B (SF1B) helicases and comprised with five domains: the zinc binding domain (ZBD), the stalk domain, domain 1B, domain 1A, and domain 2A (see Figure 1(a)).^16^ As is common among all SF1 and SF2 helicases, domains 1A and 2A, also referred as RecA-like domains, constitute the conserved helicase region. The 1A and 2A domains forms ATP binding site (ATP-pocket) at the interface as well as the RNA-cleft at the interface of domain 1B. Within 1A and 2A lie a set of highly conserved motifs essential for SF1 helicase function. These motifs include nucleoside triphosphate (NTP) binding and hydrolysis motifs (I and II), RNA binding and unwinding motifs (Ia, Ib, and IV), as well as motifs that bridge the two binding regions (III, V, and VI). Similar to other helicases, nsp13 is also a motor protein using NTP hydrolysis energy to translocate along single-stranded nucleic acid and along the way, unwinds duplex RNA or DNA from 5*′* to 3*′*.^16,17^

**Figure 1:**
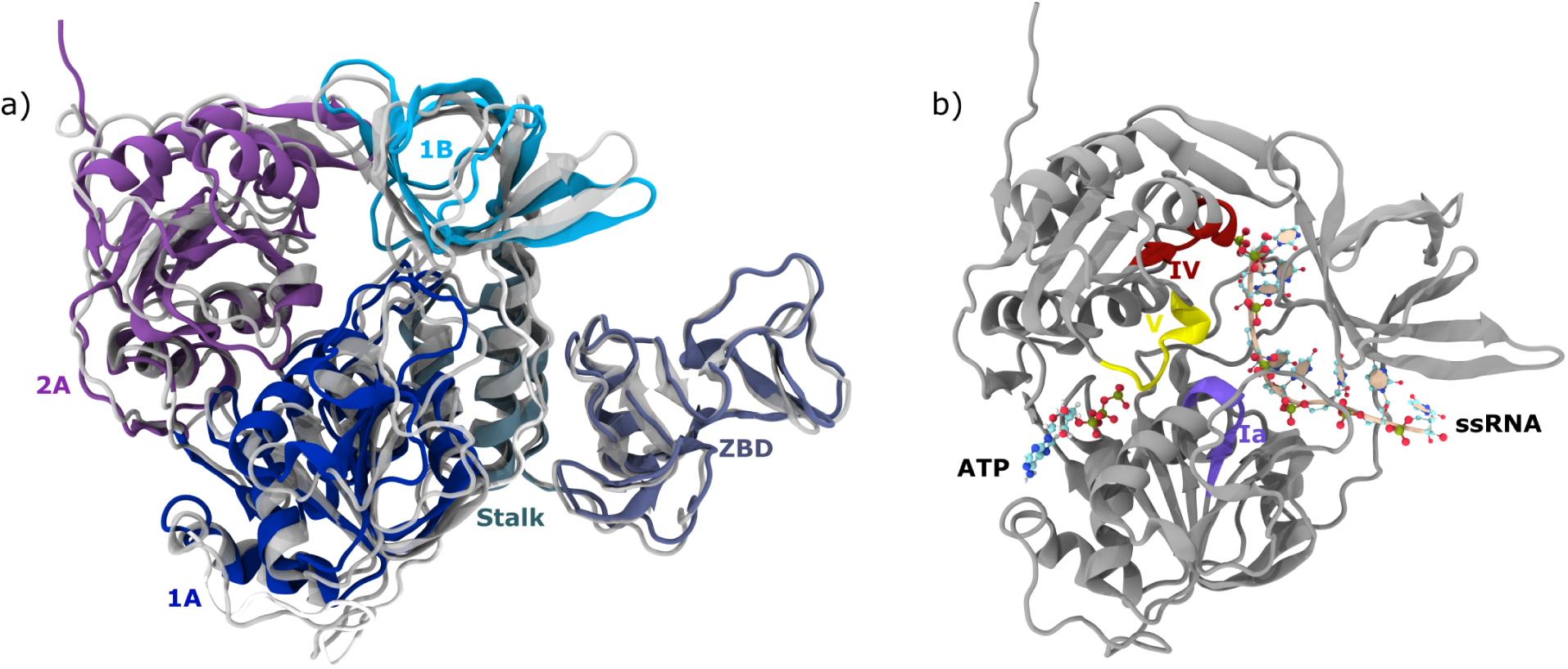
Depiction of the SARS-Cov-2 nsp13 3D structure and substrates binding sites. a) Overlay of the recent SARS-Cov-2 nsp13 structure (PDB:7NN0^9^) depicted with distinct subdomain colors and the SARS-CoV-1 nsp13 structure (PDB:6JYT^21^) in grey. The two structures have backbone RMSD of 3.06 Å. b) Depiction of the helicase domains of SARS-Cov-2 nsp13 with ATP and ssRNA bound. Important motifs (**Ia**, **IV** and **V**) of translocation are also displayed. Protein images are rendered in cartoon representation while ATP and ssRNA in ball-stick method.

The investigation of ATP hydrolysis states offers invaluable mechanistic insights into the unidirectional translocation process of SARS-CoV-2 nsp13 along single-stranded RNA (ssRNA). SF1 helicases employ two distinct mechanisms for directed translocation: the ‘inchworm stepping’ mechanism, in which two protein sites alternate in binding strength with ssRNA,^18^ and the ‘Brownian ratchet’ method, which relies on either strong or weak binding of the entire protein to ssRNA.^19^ Previous studies strongly suggest that SARS-CoV-2 nsp13 predominantly utilizes the inchworm mechanism for translocation. This process involves coordinated domain movements in a multi-step sequence to traverse a single base pair.^20–22^

A growing body of evidence outlines a series of steps that nsp13 may undergo during translocation: ATP binding-mediated transition from an ‘open’ to a ‘closed’ ATP-pocket, resulting in alterations in the binding strength of domains 1A and 2A with ssRNA; hydrolysis, triggering a rearrangement of domain affinity for ssRNA; in the product state, the ATP pocket enlarges, and the release of ADP further influences domain interactions with ssRNA. In each of these steps, ssRNA exhibits a strong interaction with either 1A or 2A. According to Jia *et al.*,^21^ the progression of one nucleotide occurs in the presence of ADP+P_i_ when the ATP-pocket is in a ‘mid-open’ conformation. This contrasts with the study by Chen *et al.*,^14^ which focuses on the ADP-bound condition where the ATP-pocket adopts an ‘open’ conformation. Moreover, evidence also indicates varying flexibility of the 1B domain during translocation^14,21^ and ambiguity in the steps of each translocation cycle.^20,21^ While these studies offer valuable mechanistic insights, they do not provide an ensemble picture of the underlying process.

The mechanism of translocation on ssRNA synergizes with the protein’s conformational ‘selection’ or ‘induction’ process. The presence of hydrolysis substrate and products at the ATP-pocket of nsp13 can be considered a mechanism for protein-ligand binding or release. Earlier, based on X-ray crystallography studies, it was believed that a protein represented a single conformation,^23^ and ligand binding *induced* the protein to undergo a new favorable conformation.^24^ This behavior is defined as the ‘conformational induction’ process. However, the conformational ensembles generated from NMR and MD simulation studies have shown that proteins are inherently dynamic and do not exist solely as a single conformation but as a thermally accessible statistical ensemble of conformations. Often, ligand binding leverages the conformational ensemble by *selecting* a preferred conformation, leading to a population redistribution of conformations. This process is termed ‘conformational selection’. In both translocation mechanisms, a conformational change in the protein is necessary to display differential affinity for ssRNA, and it can be facilitated by either conformational ‘induction’ or ‘selection’, or both processes. Studies on other motor proteins indicate that the inchworm mechanism is underlined by the *induction* effect, leading to a new conformation, while in the ratchet mechanism, substrate hydrolysis leads to forward movement by *selecting* a preferable conformation from the thermally equilibrated pre- and post-translocate states.^25,26^ Comprehensive sampling and quantifying specific structural changes can help identify the predominant underlying process of conformational change for ATP-dependent translocation in nsp13.

Previously, our group studied the translocation mechanism of SARS-CoV-2 by modeling the nsp13 structure based on SARS-CoV-1 in the pre-hydrolysis state, i.e., when nsp13 is bound to ATP and ssRNA. No significant changes were observed in the ATP-pocket due to the presence of ATP, but both the 1A and 2A domains became distant from the 1B domain. These observations differ from those of other studies where one or both of RecA domains close to 1B.^14,21^ Interestingly, the dynamics of motifs **Ia** and **IV** suggest an ‘inchworm stepping’ translocation mechanism at the apo state (i.e., when nsp13 is bound to ssRNA), and the presence of ATP leads to a probability redistribution of four states. This indicates a conformational ‘selection’ process at the RNA-cleft but is not sufficient to exclude the *inductive* effect of the hydrolysis substrate and products. Moreover, some motor proteins reveal a conformational ‘induction’ effect upon pyrophosphate release.^26^ This necessitates the investigation of the plausibility of both processes that drive the translocation mechanism in nsp13.

## Methods

### Starting Structures

In the current study, we used the crystal structure of the catalytic active form of SARS-CoV-2 nsp13 helicase, specifically chain A, from the latest crystallographic data (PDB: 7NN0^9^), resolved at a resolution of 3.04 Å. This structure was found to be co-crystallized with various elements, including substrate ANP, co-factor magnesium, water molecules, and zinc molecules within the ZBD. Additionally, ssRNA was inserted from a previously aligned RNA-bound Upf1 helicase structure (PDB: 2XZL^27^) into nsp13, as described in our previous work.^22^ Following this, we prepared four different substrate states of bound ssRNA by adding the missing residues Ala1, Asp204, Tyr205, Gly206, Asp207, Arg594, Arg595, and Asn596. These systems included ssRNA, ssRNA+ATP, ssRNA+ADP+P_i_, and ssRNA+ADP, facilitating the detailed investigation of the translocation mechanism during the hydrolysis cycle. The conversion of NH (in ANP) to an oxygen atom yielded ATP, which, after hydrolysis, formed the ADP+P_i_ complex. The release of P_i_ enabled the modeling of ADP+ssRNA, while further ADP release yielded the ssRNA system.

### System Preparation

The applied force field parameters for nsp13 and ssRNA are ff14SB^28^ and ff99bsc0*χ*OL3,^29,30^ respectively. The parameters for ATP and ADP were retrieved from the AMBER parameter database.^31^ The P_i_ parameter is available from a previous study by our group, ^32^ and the three zinc molecules of the nsp13 ZBD have been parameterized within the MCPB module of AMBER.^33^ These zinc ions form bonds with Cys-Cys-Cys-Cys, Cys-Cys-Cys-HID, and Cys-Cys-HID-HIE at their respective binding pockets. Additionally, we used the parameters of Li *et al*.^34^ for the co-factor Mg^2+^ at the active site and retained the three co-crystallized water molecules from the starting protein structure in all simulations. Each substrate state was solvated using the TIP3P water model with a 12 Å buffer, resulting in approximately 206 thousand atoms after adding Na^+^ and Cl^−^ ions to neutralize the system and mimic a 0.1 M salt concentration.

### Simulation Protocol

We employed the Gaussian accelerated molecular dynamics (GaMD) simulation technique to enhance conformational sampling in all systems. This involved applying a boost potential of 6 kcal*·*mol^−1^ on both the entire system potential (*σ*_0P_) and dihedral (*σ*_0D_). According to the protocol, GaMD simulations began from equilibrated states generated by short classical molecular dynamics (cMD). In general, both simulations were executed using the GPU-accelerated AMBER18 package^35^ with a 2 fs integration time step. Hydrogen atoms were constrained using the SHAKE algorithm.^36^ Long-range electrostatic interactions were computed using Particle Mesh Ewald (PME),^37^ with non-bonded interactions cutoff at 12 Å. Temperature and pressure were maintained using a Monte Carlo (MC) barostat and a Langevin thermostat, respectively.

The cMD simulations started with energy minimization in ten steps to gradually decrease the restraint while maintaining initial coordinates. Each run involved 2000 steps of steepest descent. In the first minimization step, a restraint of 500 kcal*·*mol^−1^ *·* Å^−2^ was applied to all non-water (excluding crystal water) and non-salt atoms. Successively, the restraint was reduced to 50.0, 10.0, 1.0, and 0.1 kcal*·*mol^−1^ *·* Å^−2^ on atoms of the protein side chain, RNA, substrate, and Mg^2+^ in four steps. The protein backbone restraint was later gradually decreased in four stages from 50.0, 10.0, 1.0, and 0.1 kcal*·*mol^−1^*·*Å^−2^ step ran without any restraint. Next, the protein complexes were gradually heated from 0 K to 300 K over 1 ns with a harmonic restraint of 40 kcal*·*mol^−1^ *·* Å^−2^ on the protein, RNA, and substrate atoms. The heating was followed by six steps of pressure equilibration to 1 atm with reducing harmonic restraints of 40.0, 20.0, 10.0, 5.0, 1.0, and 0.1 kcal*·*mol^−1^ *·* Å^−2^ on all protein and ligand atoms. The total simulation time for the first equilibration step was 1 ns, while the others were 200 ps each. Finally, we conducted a production run without restraint at constant temperature and pressure (NPT) for 10 ns in triplicate. The minimum, average, maximum, and standard deviation values of the system potential were obtained from the production run of cMD.

At the end of cMD, the GaMD simulation was initiated with equilibration for 40 ns with a boost potential. Afterward, we proceeded with the production run in three or four replicates, yielding approximately 11 *µ*s per complex, except for ADP+ssRNA, which is 16 *µ*s. Cumulatively, the ensembles amounted to approximately *∼* 50 *µ*s.

### Clustering

In this study, we used two types of clustering algorithms to identify unique conformations in substrate bound nsp13 protein: Variational Bayesian Gaussian Mixture Model (bGMM^38^) and Size-and-shape space Gaussian Mixture Model (ShapeGMM).^39^ Both models fit a multivariate dataset to a given number of Gaussian distributions to assign each data point to a cluster, but they differ in their feature sets. In bGMM, we used specific protein-protein and protein-ssRNA distances as features, whereas Shape-GMM uses the positions of ‘N’ particles as features.

In Shape-GMM, the mean and covariance are generated from the global alignment of the ssRNA system trajectory, which is used to align the other systems as a measure of translational-rotational invariance among the simulated systems. Translation is resolved by removing the center of geometry of each frame, and rotation is computed in the form of the Kronecker product of covariance, denoting ‘weighted’ alignment. After that, clustering is performed for a specified number of clusters with random frame initialization. To determine a reasonable number of clusters, the elbow heuristic combined with cross-validation (CV) is used. The elbow heuristic suggesting choosing the point along the log likelihood as a function number of clusters at which the second derivative is minimized. CV, or estimating the log likelihood of the model on data on which it was not trained, is a way to ensure the model is not overfit.

### Linear Discriminant Analysis (LDA)

LDA is a dimensionality reduction algorithm that separates clusters by minimizing intra-cluster variance while maximizing inter-cluster variances. This supervised technique uses aligned and cluster-labeled particle positions to learn cluster-separating vectors. LDA generates K-1 vectors that best separate the clusters and thus, represent a good estimate of reaction coordinate.^40^ We utilized the Python library scikit-learn with the single-value decomposition (SVD) method to handle variance.

### Model Corroboration

#### Presence of SF1 Helicase like Conserved Contacts at the ATP-pocket and RNA-cleft

In our current models, the ssRNA is incorporated from the crystal structure of Upf1 helicase,^27^ and all the substrates are generated by subtle modifications of ANP, which is co-crystallized with SARS-CoV-2 nsp13.^9^ Due to such alterations, it is important to check whether these insertions are stable by corroborating protein contacts with crystal structures of other SF1 helicases. First, we identified protein residues within 5.0 Å of ATP in Upf1^27^ and IGHBMP helicase^41^ and computed the occurrence probability of similar contacts in the simulations of systems. A contact is considered whenever a protein residue is present within 5.0 Å of ATP, ADP+P_i_, and ADP. The probability of contacts at the RNA-cleft and ATP-pocket is presented in Table S1 and Table S2, respectively, by considering residues of important motifs. A greater number of contacts show persistence of 65-100% in all substrate systems, although very few contacts at the RNA-cleft show substrate specificity. These analyses suggest effective model preparation for all substrates.

#### Comparing Conformational Ensemble of Pre-hydrolysis states of SARS-CoV-2 nsp13 current Model with Previous Modelling

Previously, our group modeled SARS-CoV-2 nsp13 based on the SARS-CoV-1 crystal structure, and analyses of pre-hydrolysis systems suggested a four-state model of the inchworm translocation mechanism.^22^ These states are identified through clustering based on the distances of motifs **Ia** and **IV** to the nearest phosphates (see Figure 1(b) for these motifs location), as these motifs play an important role in nucleic acid unwinding. ^42,43^ The residue identities of these motifs for distance calculation are provided in Table S3. To compare between the two models, we investigated whether similar states were observed in the ssRNA and ssRNA+ATP substrate states of the new model. To do so, we used the same bGMM and LDA procedure from our previous work. The results of distance projection onto LD1 and LD2 vectors are shown in Figure S1. Consistent with our previous simulations, LD1 separates S4, while LD2 separates S1, S2, and S3. We computed the population of states in the current model simulation (see Table S5) and noticed that S1 is present in a minuscule proportion, but a similar trend of population redistribution is observed in the ATP+ssRNA-bound nsp13 system.^22^ Altogether, these analyses are qualitatively comparable with our earlier results.

## Results and Discussion

Understanding of the ssRNA translocation mechanism relies on identifying crucial structural and conformational changes in nsp13 that occur in different states of the ATP hydrolysis cycle. In this study, we investigate the possibility of both the conformational ‘selection’ and ‘induction’ processes in the ATP-dependent nsp13 translocation cycle. To achieve this, we analyze simulations of ssRNA, ssRNA+ATP, ssRNA+ADP+P_i_, and ssRNA+ADP-bound nsp13 helicase models as representations of hydrolysis states. We initiate our discussion by evaluating both the ‘selection’ and ‘induction’ processes for the global motion of protein domains (1B, 1A, 2A), followed by an examination of local motion at the RNA cleft. Based on these insights, we propose a plausible mechanism of translocation

### ATP-dependent Protein Motion Demonstrates both Conformational ‘Selection’ and ‘Induction’ Process

Identifying ATP-dependent protein conformations unveils the underlying mechanism of protein motion. Experimental and molecular simulation studies have highlighted significant changes in 1A and 2A upon ATP binding, as well as movement of 1B towards either 1A or 2A, depending on the translocation stage.^14,21^ However, all these insights are based on specific hydrolysis substrates and are limited in providing a comparative description of conformational changes between the hydrolysis states. In this study, we investigated protein motion and explored the possibilities of conformational ‘selection’ and ‘induction’ using weighted ShapeGMM (W-SGMM) based clustering at the C*_α_* resolution of domains 1B, 1A, and 2A. Our approach involved evaluating these processes by amalgamating the trajectories of all simulated systems to create a single trajectory, followed by clustering analysis. We hypothesized that if conformational ‘induction’ exists along the hydrolysis cycle, the clustering would reveal unique clusters or conformations specific to each hydrolysis substrate.

The ATP-dependent protein motion of nsp13 indicates the presence of four unique conformations. A plot of log-likelihood per frame as a function of cluster numbers for the training set (blue) and CV set (orange) is presented in Figure 2(a). The CV set significantly diverges from the training set after four clusters, which suggests overfitting of our model with more than four clusters. The second derivative of log-likelihood as function of clusters on the training set is minimized at n_c_lusters = 4. Combined, these data suggest that the protein ensemble is well described by four conformational clusters. We denote these clusters as ^P^C1, ^P^C2, ^P^C3, and ^P^C4. To demonstrate the distinguishability of these conformational clusters, we performed linear discriminant analysis on these clusters and projected the trajectory of C*_α_* atoms onto the resulting LD1 and LD2 vectors, as presented in Figure 2(b). LD1 separates ^p^C2 from other clusters while LD2 and LD3 separates ^P^C1, ^P^C2, and ^P^C4.

**Figure 2:**
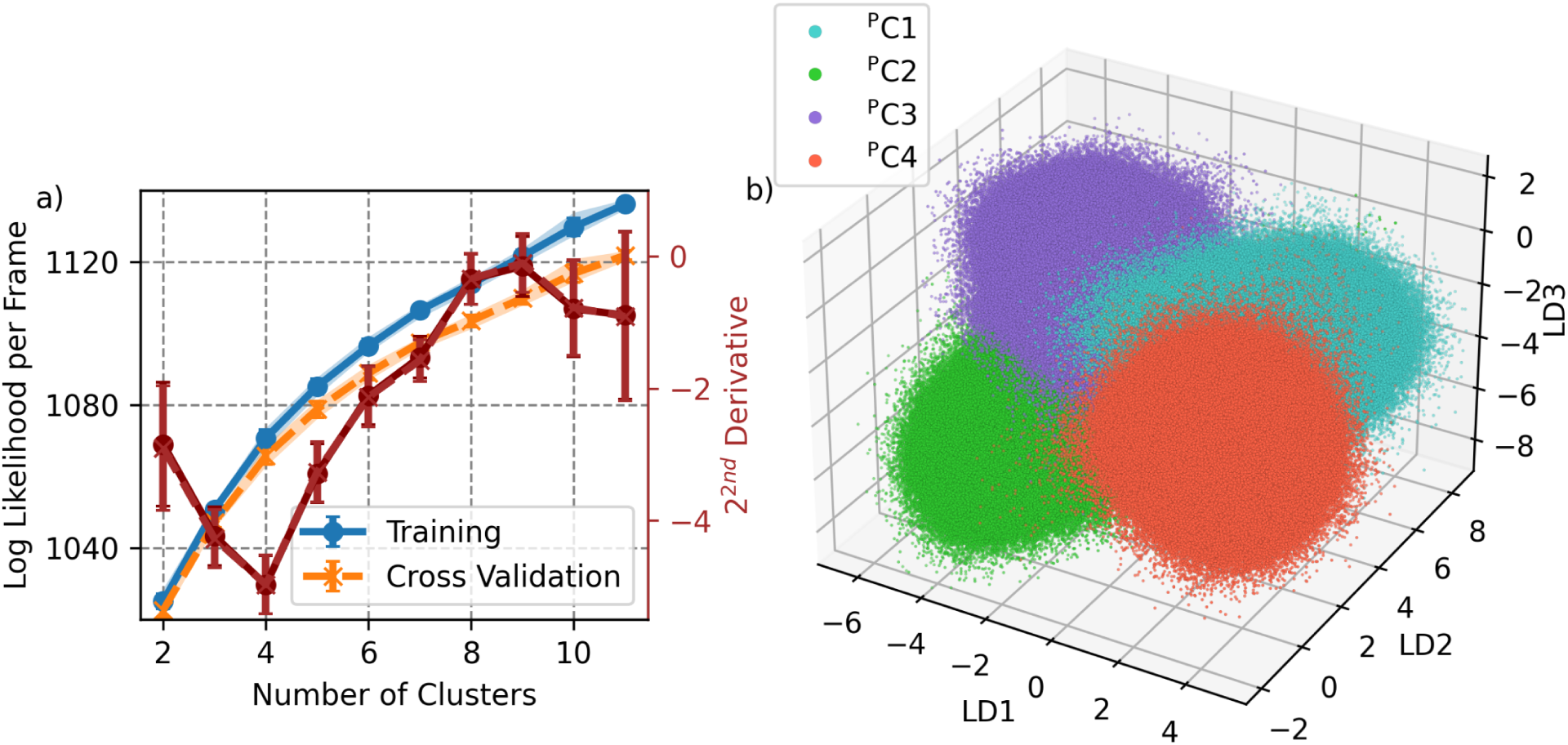
Protein conformational states of nsp13 from the combined ssRNA, ATP+ssRNA, ADP+P_i_+ssRNA, and ADP+ssRNA trajectories determined by shapeGMM. a) The log likelihood per frame as a function of number of clusters is plotted for training (blue) and CV (orange) sets computed by W-SGMM method. We used 80k and *∼*331k frames for training and CV set respectively. The error bars denote standard deviation of sampling 5 different training set and shaded region marks 90% confidence interval.The second derivative of log-likelihood as a function of cluster number is plotted in red with values given on the right-hand y-axis. b) This 3D plot denote the projection of amalgamated trajectory comprises C*_α_* atoms of 1B, 1A and 2A domains onto LD1, LD2 and LD3 eigenvectors. Each protein cluster is marked with a unique color, see legend.

Sampling of protein cluster exhibits a directional correlation with hydrolysis progression. Figure 3 displays the probability distribution of protein clusters for each system. In the ssRNA and ssRNA+ADP systems, the sampled clusters were ^P^C1, ^P^C2, and ^P^C4, while the ssRNA+ATP and ssRNA+ADP+P_i_ systems predominantly sampled ^P^C1, ^P^C2, and ^P^C3 clusters. This suggests hydrolysis substrate and products dependent protein motion. In a long, unbiased simulation, Matteo *et al.*^44^ also reported two conformations in the ssRNA system, indicating conformational variation. Furthermore, the prevalence of ^P^C1, ^P^C2, ^P^C3, and ^P^C4 clusters in ssRNA, ssRNA+ATP, ssRNA+ADP+P_i_, and ssRNA+ADP-bound nsp13, respectively, hints at the directionality of the hydrolysis cycle. This sequence suggests that the cycle might commence with the population of ^P^C1, followed by a predominant sampling of ^P^C2, then transitioning to ^P^C3, and ultimately concluding with the population of the ^P^C4 cluster.

**Figure 3:**
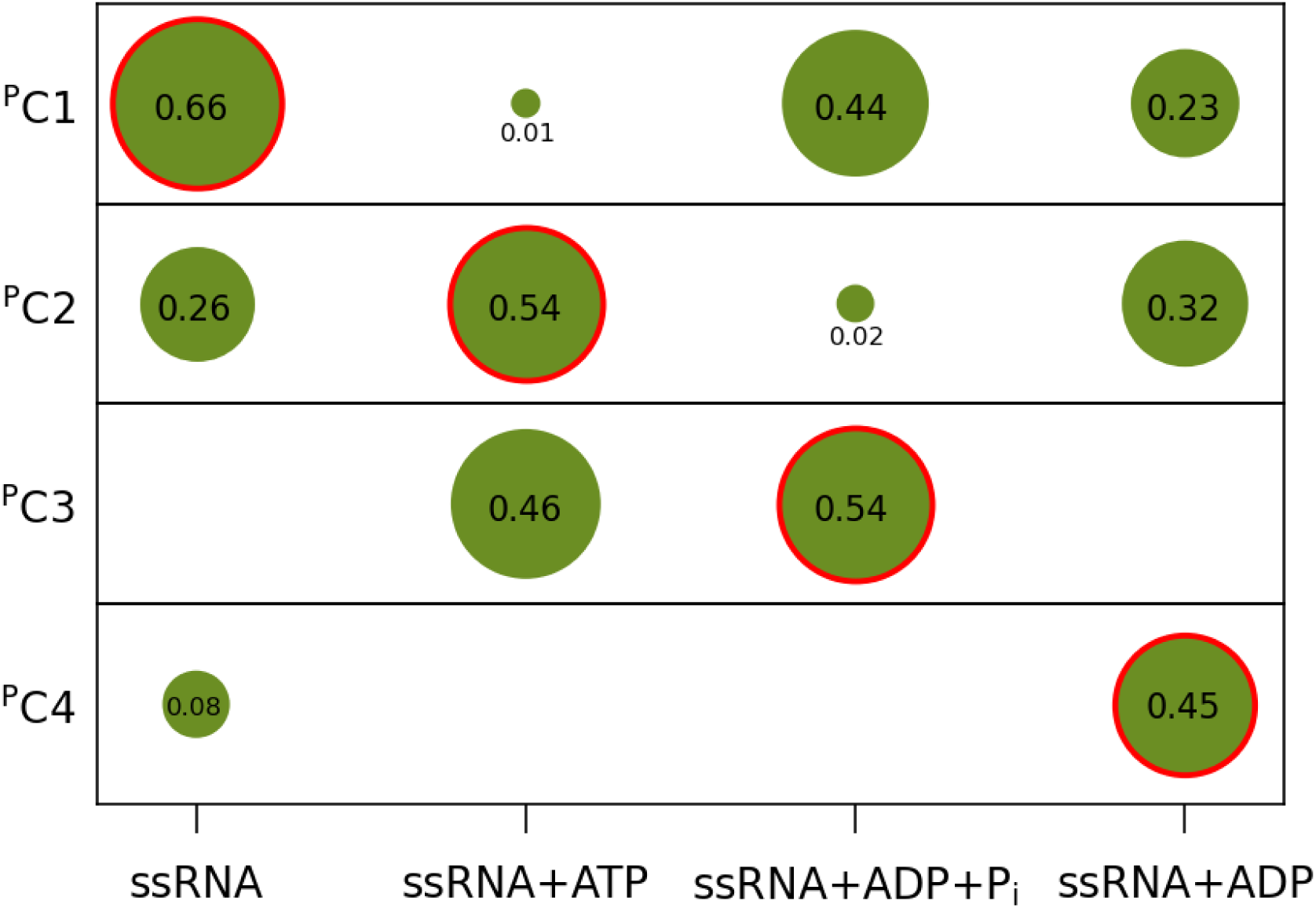
Hydrolysis substrate directed sampling of protein states. The probability of states explored by each system is plotted here and presented in scatter form. The size of each scatter is proportional to the respective states probability. The states are identified by ShapeGMM clustering on C*_α_* position of 1B, 1A and 2A domains of amalgamated trajectory.

The sampling of ATP-dependent protein conformations underscores the significance of both conformational ‘selection’ and ‘induction’ mechanisms. It is noteworthy that all systems sample ^P^C1 and ^P^C2 clusters, yet the populations of these clusters vary significantly among the hydrolysis states. This distinctive feature is indicative of the ‘selection’ process, where the presence of hydrolysis substrate (ATP) influences the ‘selection’ of a specific conformation from the existing conformational ensemble, resulting in a redistribution of populations. In addition to the ‘selection’ process, ATP binding initiates the sampling of the ^P^C3 cluster, which can be regarded as a novel conformation compared to the ssRNA-bound state. This highlights that the presence of ATP *induces* nsp13 to adopt a new conformation as part of the ‘induction’ process. In the ADP+P_i_ bound condition, nsp13 samples all the existing clusters observed in the ATP-bound state but predominantly *selects* the ^P^C3 conformation. Conversely, in the presence of ADP, nsp13 predominantly samples the ^P^C4 cluster but lacks the ^P^C3 cluster. This indicates that the absence of P_i_ *induces* nsp13 to explore a new conformation. Consistent with these findings, Chen *et al.* also noted in their study that ADP-bound nsp13 explores a unique conformation compared to ATP-bound nsp13.^11^ In the ssRNA-bound state, nsp13 explores all the existing conformations observed in ADP-bound nsp13 but shows a preference for *selecting* ^P^C1. Collectively, these observations suggest that protein motion is influenced by both conformational ‘selection’ and ‘induction’ processes.

### Protein Clusters Demonstrate Nucleotide Binding, Release and Translocation Mechanisms

The characterization of protein clusters describes the motion of protein domains (1B, 1A, and 2A) in relation to the hydrolysis cycle and translocation. To characterize these clusters, we calculated angles between the vectors representing the center of mass distances between domains. A graphical representation of these angles on the nsp13 structure is presented in Figure 4. The angle *θ*_1_ measures the status of the 1A-2A domain cleft and is defined by the vectors (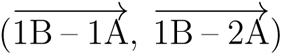). Similarly, *θ*_2_ and *θ*_3_ are designated for the 1A-1B and 2A-1B clefts, respectively, and are defined by the vectors (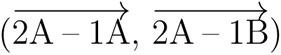) and (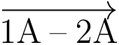, 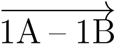), respectively. Each angle value reflects the motion of all three domains (1B, 1A, and 2A) and can therefore be interpreted as the global motion of nsp13.

**Figure 4:**
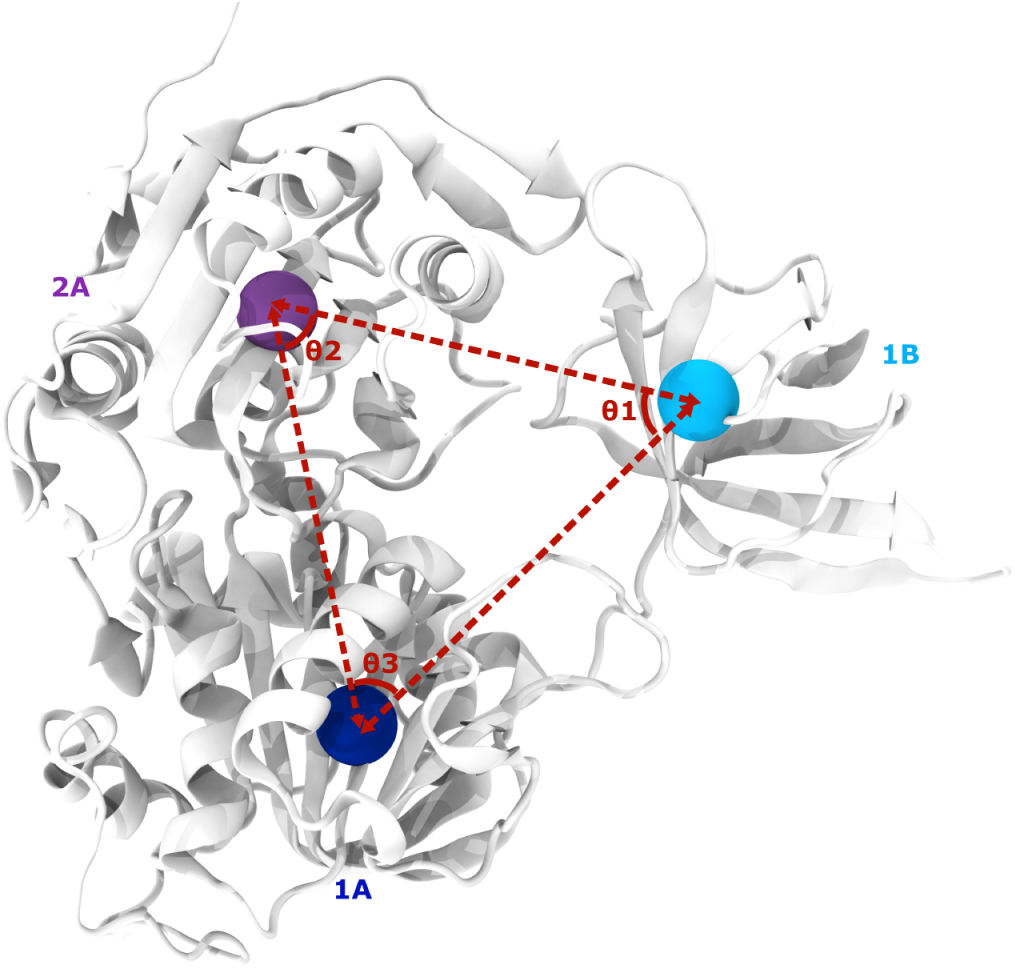
Depiction of inter-domain angles within the nsp13 structure. The spheres represents center-of-mass location and colored to its respective domain.

Protein clusters leads transition from ‘open’ to ‘closed’ or *vice versa* of the 1A-2A, 1A-1B, and 2A-1B clefts, mediated by ‘mid-open’ conformation. The angle values for each cluster are provided in Table 1. The largest (smallest) average value of any angle is labeled as the ‘open’ (‘closed’) conformation, while a value near the median of these extremes is denoted as the ‘mid-open’ conformation. Representative structures for each conformation are shown in Figure 5. We attempted to interpret the transitioning mechanism of the four protein clusters based on these angle values. We initiated from ^P^C1, as indicated by substrate-specific protein cluster probabilities:

- In the first step, 1A moves downward while also moving towards 1B, resulting in all clefts becoming ‘mid-open’. This leads the protein into ^P^C2.
- In the second step, 1B moves downward. While the 1A-2A cleft remains in the ‘mid-open’ conformation, the 1A-1B and 2A-1B clefts switch to ‘closed’ and ‘open’ conformation, respectively. This transition progresses the protein into ^P^C3.
- During the third step, 2A move downward by rotating towards 1B, resulting in an ‘open’ 1A-2A cleft, while the other two clefts remain ‘closed’. This causes the protein to shift into ^P^C4.
- In the fourth step, the 1A domain rotates towards 2A, resulting in ‘closed’, ‘open,’ and ‘closed’ clefts for 1A-2A, 1A-1B, and 2A-1B, respectively. Subsequently, the protein re-enters ^P^C1.
- In this cycle of four protein clusters, the 1A-2A and 2A-1B clefts undergo a ‘closed’ to ‘open’ transition, while the 1A-1B cleft moves from ‘open’ to ‘closed’. The presence of ^P^C2 signifies the ‘mid-open’ conformation as an intermediate state during cleft transitions.

**Figure 5:**
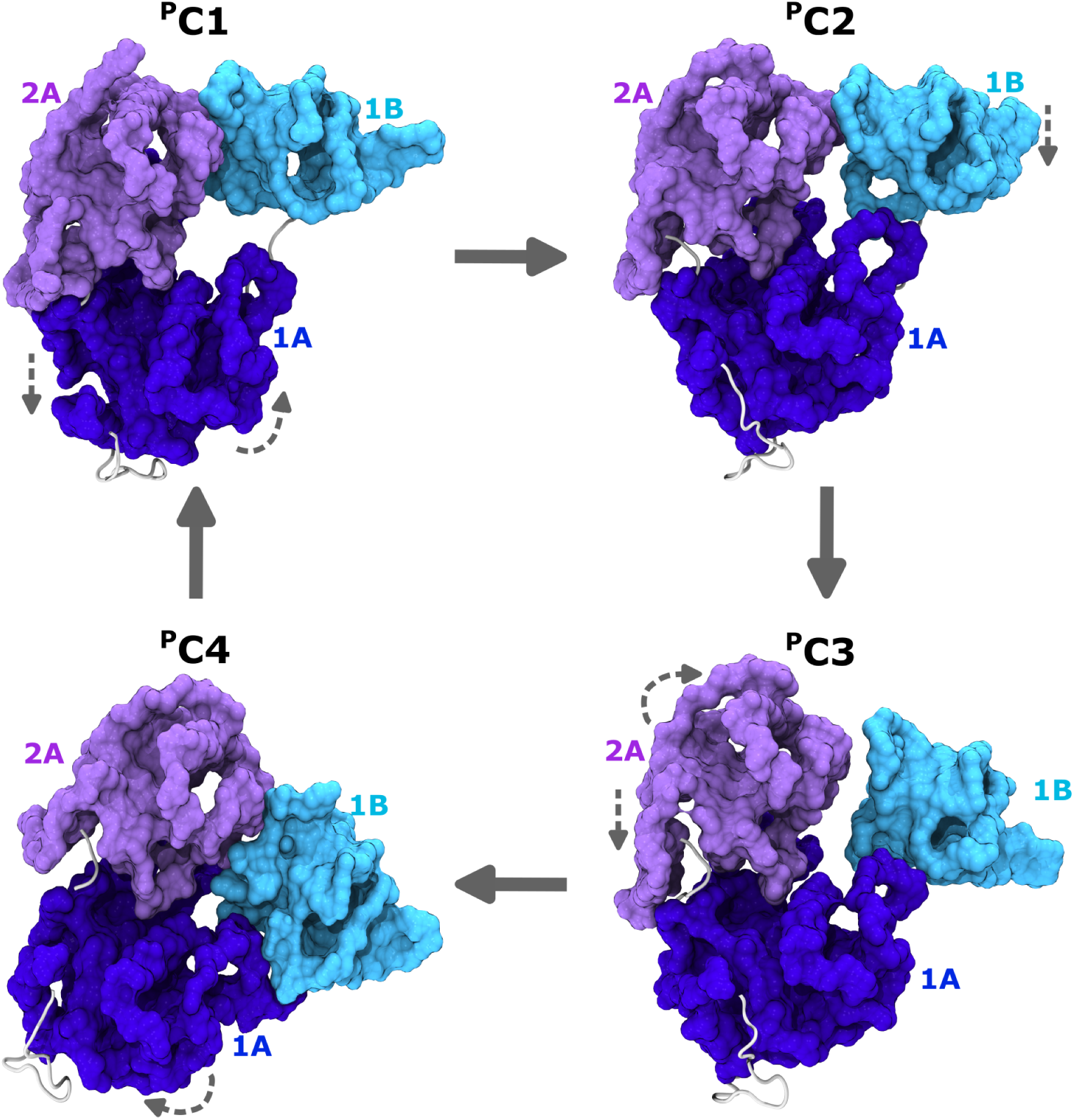
Representative structures of protein clusters comprising 1A, 2A and 1B domains. These clusters are identified in W-SGMM protein clustering and arranged as a four-state cycle based on substrate-specific sampling. The broken arrow indicate domain motion for transitioning between protein clusters as directed by solid arrow. The domains are rendered in ‘surf’ mode of VMD.

**Table 1:**
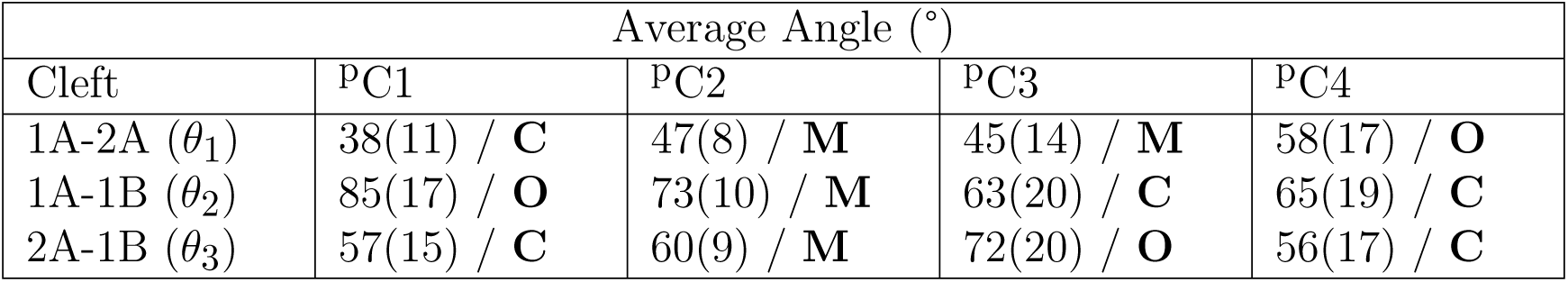
Average (standard deviation) values of angles of inter-domain clefts for each protein clusters. Each cleft marked as ‘Open’ (O), ‘Closed’ (C) and ‘Mid-open’ (M) based on the minimum, maximum and medium values of respective angle of protein clusters.

| Cleft | Average Angle (°) |  |  |  |
| --- | --- | --- | --- | --- |
|  | PC1 | PC2 | PC3 | PC4 |
| 1A-2A ($\theta_1$) | 38(11) / **C** | 47(8) / **M** | 45(14) / **M** | 58(17) / **O** |
| 1A-1B ($\theta_2$) | 85(17) / **O** | 73(10) / **M** | 63(20) / **C** | 65(19) / **C** |
| 2A-1B ($\theta_3$) | 57(15) / **C** | 60(9) / **M** | 72(20) / **O** | 56(17) / **C** |

ATP-dependent sampling indicates the binding and release of nucleotide. In absence of nucleotide in the ATP-pocket, the 1A-2A cleft adopts a ‘closed’ or ‘mid-open’ conformation, ssRNA bound nsp13 mostly sampled ^P^C1 and ^P^C2, with ^P^C1 being predominant. The ‘closed’ conformation aligns with the findings of Chen *et al.* in their nsp13_t_-engaged conformation.^14^ However, ATP binding *selects* the ‘mid-open’ conformation of the 1A-2A cleft, as observed in the ssRNA+ATP system, where ^P^C2 and ^P^C3 conformations are predominantly sampled. This observation contrasts with other studies which reported ‘closed’ 1A-2A cleft.^14,21^ This deviation can be attributed to the use of different metrics and strict cutoffs for defining the cleft status.

Interestingly, ADP+P_i_ bound nsp13 *selects* both ^P^C1 and ^P^C3 equally. In these clusters, the 1A-2A cleft explores both the ‘closed’ and ‘mid-open’ conformations, suggesting a potential mechanism by which nsp13 guards against product release through this cleft. However, the 2A-1B cleft samples ‘closed’ and ‘open’ states, with the ‘open’ 2A-1B cleft also observed in ATP-bound nsp13. We propose that the ‘open’ 2A-1B cleft in ^P^C3 may be specific to P*_γ_* or P_i_ and could facilitate P_i_ release. In the presence of ADP alone, nsp13 explores an ‘open’ 1A-2A cleft, with ^P^C4 being the most sampled conformation, along with ‘closed’ (^P^C1) and ‘mid-open’ (^P^C2) conformations to some extent. The ‘opening’ of the 1A-2A cleft aids in ADP release and may be indicative of the *induction* of P_i_ release. ^P^C4 like conformation is also reported by Chen et al.^14^ when ADP is present in the ATP-pocket. In summary, the ‘mid-open’ and ‘open’ 1A-2A cleft is involved in ATP binding and ADP release respectively, whereas the ‘open’ 2A-1B clefts mark a probable release pathway for P_i_.

The transitioning mechanism between protein clusters indicates a downward motion of domains, representing translocation from 5*′* to 3*′* of the ssRNA tract. We observe a downward motion of the 1A domain during the transition from ^P^C1 to ^P^C2. Both of these clusters are present in the ssRNA system, but ^P^C1 is more sampled than ^P^C2. However, the presence of ATP redistributes the probability, with ^P^C2 being sampled more than ^P^C1. This demonstrates that in the absence of ATP, the 1A domain fluctuates between upward and downward motion, but ATP binding stabilizes the downward motion, as indicated by the conformational ‘selection’. Our observation differs from the findings of Perez-Lemus et al.^7^ and Jia *et al.*,^21^ which suggest that ADP and ADP+P_i_ presence, respectively, induce 1A downward motion.

The 1B domain moves downward when transitioning from ^P^C2 to ^P^C3, and both of these clusters are equally and predominantly sampled in ATP-bound nsp13, indicating an *inductive* mechanism. Importantly, ADP+P_i_ bound nsp13 only samples ^P^C3, indicating the stabilization of the downward state of 1B through the *selection* process. In the case of the 2A domain, the downward motion is only observed during the transition from ^P^C3 to ^P^C4. Both of these protein clusters are sampled in the presence of ADP+P_i_ and ADP, respectively, emphasizing the *inductive* effect of the absence (or release) of P_i_. We hypothesize that the 1A motion during the ^P^C4 to ^P^C1 transition is ADP-dependent reorganization toward the 2A domain. A similar observation is also reported by Matteo *et al.*,^44^ suggesting that RNA pushes 1A toward the 2A domain. These insights depict an ATP-dependent translocation mechanism explained by the downward motion of each domain.

### ATP-dependent RNA-cleft Motion Reveals Conformational ‘Selection’ Process

In the previous section, we examined the downward motion of the 1A, 2A, and 1B domains within the protein clusters, influenced by both conformational ‘selection’ and ‘induction’ processes. However, this analysis did not delve into their motion in relation to ssRNA. Studies on SF1 helicases suggest that motifs **Ia** and **IV** play a crucial role in translocation.^22,42,43^ But clustering of distances between RNA phosphate and motifs (**Ia** and **IV**) biased toward the ‘conformation selection’ process along with limitation in properly describing the translocation mechanism. In this regard, we examined RNA cleft motion to assess the possibility of both conformational ‘selection’ and ‘induction’ processes. We achieved this by performing W-SGMM clustering on the positions of C*_α_* atoms of motifs **Ia**, **IV**, **V** and phosphate atoms of ssRNA (see Figure 1(b)), applying a similar protocol to the protein clustering. The inclusion of motif **V** helps overcome the feature limitation and studies have also emphasized its importance at the RNA-cleft.^21^ For motif residue identities, refer to Table S3, which were used in clustering.

Clustering of RNA-cleft motion identified three unique clusters. The results of a CV scan, performed on the amalgamated trajectory for selective atoms, are provided in Figure 6(a). A plot of the log-likelihood per frame as a function of the number of clusters shows that both the trained data (in blue) and the CV data (in orange) are superimposed, indicating a good fit of the model. The log-likelihood per frame sharply increases from cluster number two to seven, after which it plateaus for larger clusters. The second derivative of the log-likelihood per frame per cluster shows the lowest value at cluster number three, marking a significant change in slope. Based on these results, we clustered RNA-cleft motion, comprising selective motifs positions, into three clusters (referred to as ^R^Cs). Later, LDA analysis was performed to evaluate the distinction of these multivariate clusters in lower dimensions. In Figure 6(b), the projection of the amalgamated trajectory onto LD1 and LD2 vectors is displayed. As observed in the protein clusters, the LD1 vector separates ^R^C1, whereas the LD2 vector separates ^R^C2 and ^R^C3.

**Figure 6:**
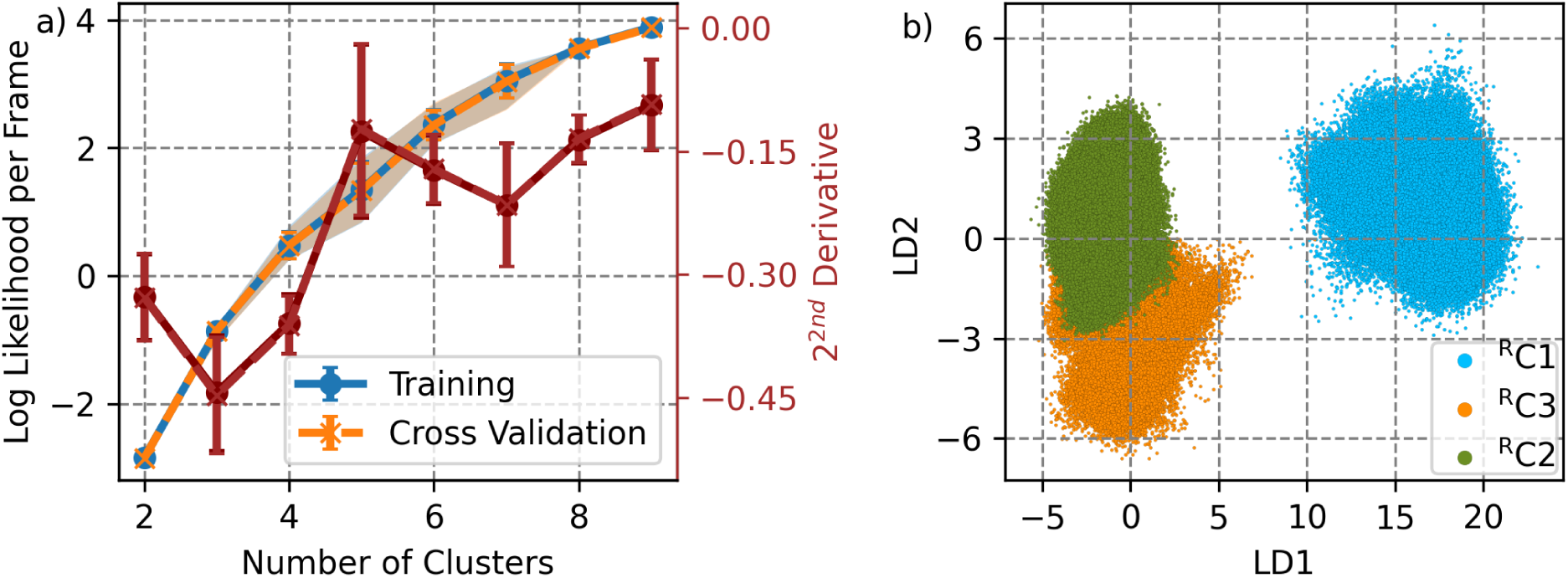
Protein-RNA conformational states determined by shapeGMM clustering. (a) The log likelihood per frame as a function of number of clusters is plotted for training (blue) and CV (orange) set. It is computed by W-SGMM method and used 80K and *∼*331K frames for training and CV set respectively. The error bar denotes standard deviation of sampling 5 different training set and shaded region marks 90% confidence interval. (b) This plot denote change of slope as a function of number of clusters. For details of this calculation, see caption of Figure 2.

Hydrolysis substrates and products redistribute the sampling of RNA-cleft clusters. We quantified cluster probabilities in each system, which are provided in Table 2. The ssRNA system sampled ^R^C1 in a very small proportion, but ^R^C2 and ^R^C3 were sampled largely. Among all the states, ^R^C2 predominates. Interestingly, the sampling of ^R^C1 is not observed for any other systems, but the sampling of ^R^C2 increases compared to the ssRNA system. Moreover, in the presence of ATP, the sampling of ^R^C2 increases, but the sampling of ^R^C3 significantly reduces. A similar trend is also observed when nsp13 is bound to ADP+P_i_. In contrast, the presence of ADP leads to reduced sampling of ^R^C2, but importantly, the sampling of ^R^C3 increases. In general, we do not observe the occurrence of hydrolysis substrate or products specific clusters for RNA-cleft motion. Instead, the probability of the ssRNA system exploring clusters is altered along the hydrolysis progression. This suggests that ATP-dependent RNA-cleft motion *selects* conformations from the existing conformational palettes along the hydrolysis cycle.

**Table 2:** Probability of states for RNA-cleft motion explored by each systems. The states identified by Shape-GMM clustering on amalgamated trajectory of motif Ia, IV, V and ssRNA phosphates.

| States | Probability |  |  |  |
| --- | --- | --- | --- | --- |
|  | ssRNA | ssRNA+ATP | ssRNA+ADP+P<sub>i</sub> | ssRNA+ADP |
| $R_{C1}$ | 0.05 | — | — | — |
| $R_{C2}$ | 0.64 | 0.94 | 0.98 | 0.84 |
| $R_{C3}$ | 0.31 | 0.06 | 0.02 | 0.16 |

### RNA-cleft Motion Indicate ‘Inchworm’ Translocation Mechanism

Identifying structural changes in RNA-cleft clusters will delineate the altered ssRNA affinity. Conformational heterogeneity at the RNA cleft suggests an underlying ‘selection’ mechanism but needs to identify structural changes. We computed the distances between motifs and the nearest phosphates to measure structural change between the conformations, which are tabulated in Table 3. These distances between atom positions can be considered as local motion at the RNA-cleft. Depending on the distances, ^R^C1 can be designated as the ‘loose’ conformation, where motifs **IV** and **Ia** are largely distant from the phosphate. In ^R^C2, all motifs are close to the phosphate and hence denoted as the ‘compact’ conformation. However, in ^R^C3, only motif **Ia** appears distant from the phosphate, and we denote this state as ‘medium-loose’. A representative structural image of each cluster is presented in Figure 7.

**Figure 7:**
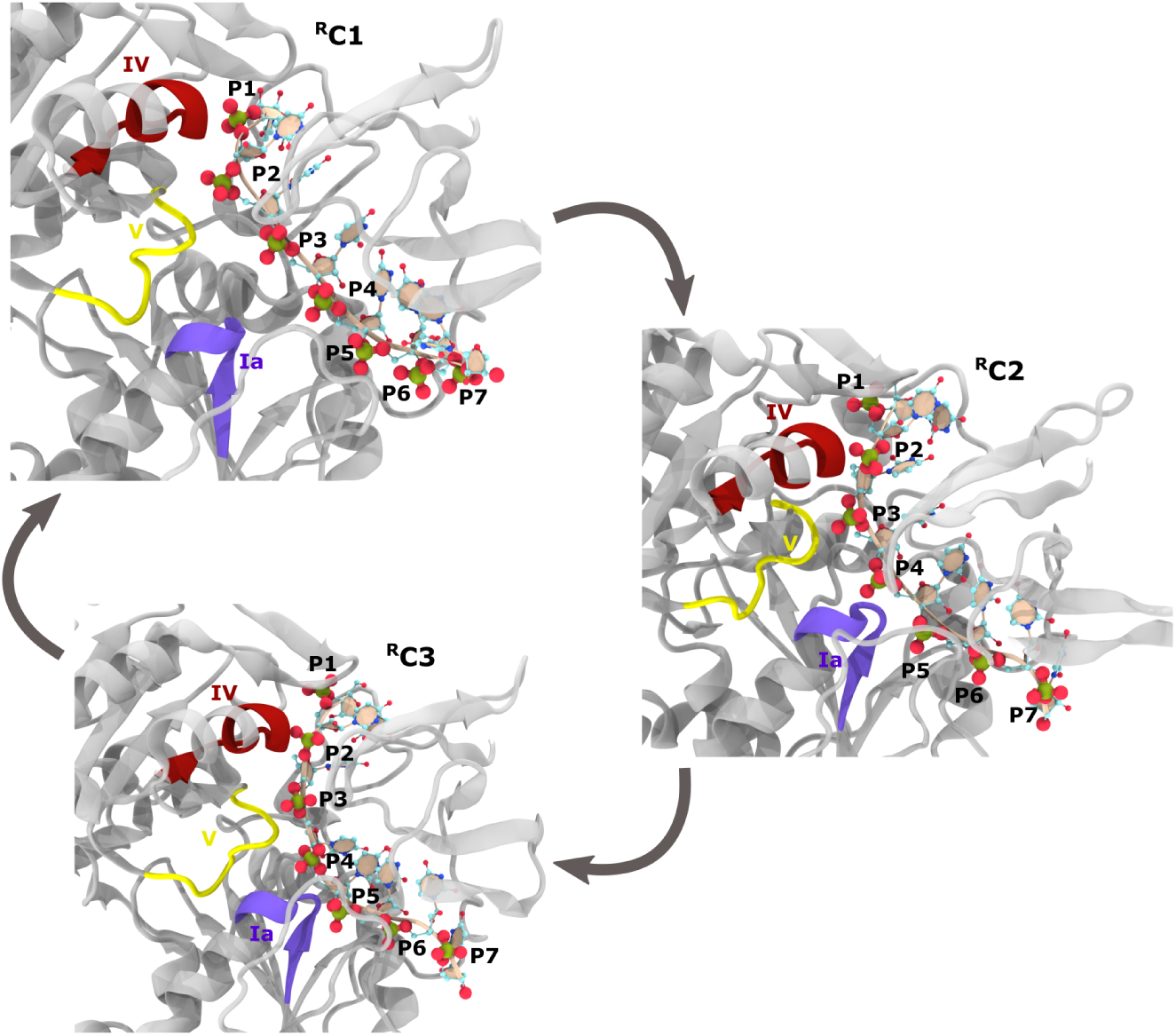
Representative structures of RNA-cleft clusters depicting motif Ia, IV, V location relative to ssRNA. Three RNA-cleft clusters are identified in W-SGMM clustering. Protein and ssRNA rendered in ‘new-cartoon’ representation of VMD. In protein structure, only motifs are colored in red, yellow and purple for motif **Ia**, **IV**, **V** respectively while remaining region colored in grey. In ssRNA, phosphate moiety showed in ‘CPK’ mode where green and red color used for phosphate and oxygen respectively.

**Table 3:** Average separation distance (standard deviation) between motif IV, Ia, V and nearest ssRNA phosphates (P) for clusters ^R^C1, ^R^C2 **and** ^R^C3.

| Average Distance (Å) |  |  |  |
| --- | --- | --- | --- |
| Residues | $R_{C1}$ | $R_{C2}$ | $R_{C3}$ |
| IV-P | 7.5(5) | 5.0(6) | 5(1) |
| V-P | 5.1(6) | 6(2) | 6(1) |
| Ia-P | 9.1(8) | 6(1) | 7(3) |

Structural changes in RNA-cleft clusters demonstrate the traversal of one phosphate through the combined motion of motifs **IV**, **V**, and **Ia**. Based on the identity of the nearest phosphate (see Table 4) for motifs in each cluster, we attempted to envisage the transitioning mechanism. Let us assume that the cycle starts from ^R^C1, the ‘loose’ conformation:

- In the first step, motif **IV** and **Ia** form a strong bound with P_n_ (P2) and P_n+2_ (P4), respectively, while motif **V** undergoes a power-stroke motion to switch its strong coordination from P_n_ (P2) to P_n+1_ (P3). After this, the RNA-cleft transitions to ^R^C2 and becomes a ‘compact’ conformation.
- In the second step, motif **Ia** detaches from P_n+2_ (P4) to form contact with P_n+3_ (P5), resulting in a ‘medium-loose’ RNA-cleft, and the protein enters ^R^C3.
- In the third step, motifs **IV** and **Ia** detach from P_n_ (P2) and P_n+2_ (P4), respectively, and become loosely bound to P_n+1_ and P_n+3_, respectively. Following the third step, the protein re-enters ^R^C1 and begin next translocation cycle.

**Table 4:** Motif Specific Closest Phosphates Probability in each clusters. The normalization performed for each motif for each clusters.

| Probability |  |  |  |
| --- | --- | --- | --- |
| Residues | $R_{C1}$ | $R_{C2}$ | $R_{C3}$ |
| IV-P2 | 100% | 94.5% | 87% |
| IV-P3 | - - | 5.5% | 13% |
| V-P2 | 97.7% | - - | - - |
| V-P3 | 2.3% | 91.7% | 89% |
| V-P4 | - - | 8.3% | 11% |
| Ia-P3 | 2.0% | - - | - - |
| Ia-P4 | 96.5% | 63.9% | 61% |
| Ia-P5 | 1.5% | 36.1% | 39% |

Throughout the entire cycle, all motifs translocate one phosphate from 5*′* to 3*′*, while motifs **IV** and **Ia** always maintain a one-phosphate gap. Interestingly, motif **V** always showed a strong contact with phosphate, even during the loose binding of both motifs **IV** and **Ia** to its nearest phosphate (^R^C1). The strong binding of motif **V** also agrees with Jia *et al.* report on SARS-CoV-1 nsp13; the H/D exchange data indicate less deuterium exchange for peptides 511 to 542.^21^ The local motions of motifs illustrate that translocation of one phosphate occurs through the loose and tight binding of motifs **IV** and **Ia**, but motif **V** always holds onto the RNA, signifying an ‘inchworm’ stepping mechanism of translocation.

### Overlap of Protein and RNA-cleft Clusters

Due to the difference in the number of particles for protein and RNA-cleft clustering, we compared two types of clusters, as shown in the matching wheel plot in Figure 8. It illustrates good overlap between protein and RNA-cleft clusters. ^P^C1, ^P^C2, and ^P^C3 include two RNAcleft clusters, namely ^R^C2 and ^R^C3, while ^P^C4 encompasses all three RNA-cleft clusters. From ^P^C1 to ^P^C3 clusters, we observed a downward motion of the 1B and 1A domains. We propose that as the RNA cleft fluctuates between ‘compact’ and ‘mid-loose’ conformations, the 1A and 1B domains move downward. In ^P^S4, we observed the 2A domain moving down, causing the RNA-cleft to transition to the ‘loose’ conformation and fluctuate between three clusters. Overall, this comparison suggests that each protein cluster is a conformational ensemble of RNA-cleft clusters (or *vice versa*), and the downward motion of domains (1B, 1A, and 2A) leads to the redistribution of RNA-cleft clusters by *selecting* the preferred state. However, the domain motion along the 5*^′^* to 3*^′^* direction is influenced by both ‘selection’ and ‘induction’ processes. The superimposition of both local and global motions results in the translocation of one phosphate for each hydrolysis turnover.

**Figure 8:**
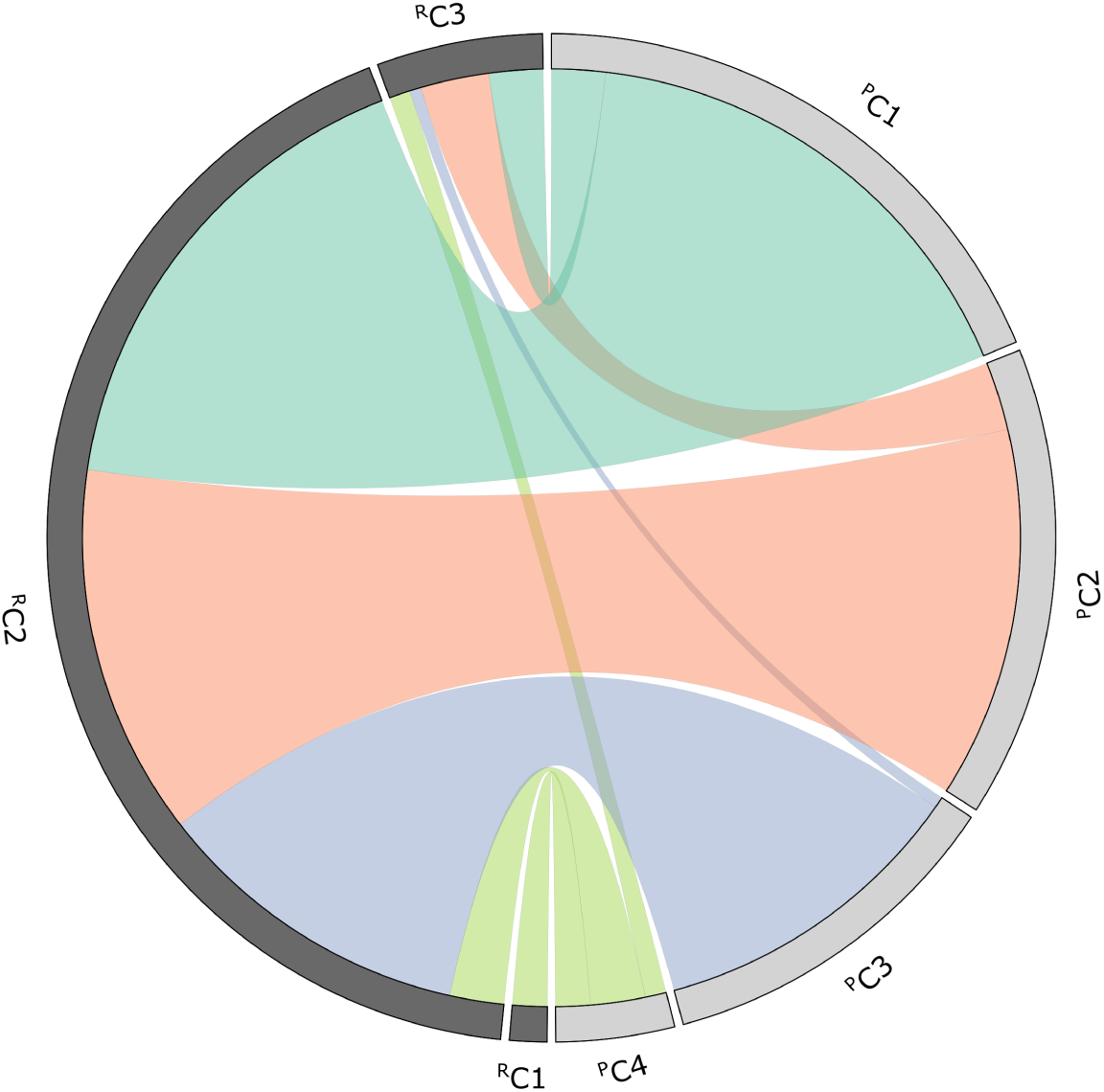
Comparison of protein and RNA-cleft states. The probability of RNA-cleft states (^R^C) in each protein state (^P^C) is computed from amalgamated labelled trajectory and plotted as matching wheel.

### Proposed Mutation to Impair the Translocation

The coordinated motion of motifs **VI**, **V**, and **Ia** implies a potential translocation of a phosphate group on the ssRNA tract, driven by the process of conformational ‘selection’. In contrast, the protein’s motion exhibits a downward shift of domains from the 5*′* to the 3*′* end, influenced by both the ‘selection’ and ‘induction’ processes. The LD eigenvectors serves as a discriminator for protein and RNA-cleft motion encapsulated clusters and constitute critical positions of each particle to separate its motion with the shapeGMM clusters. ^40^ Consequently, we computed the magnitude of the LD1 vector for the protein’s C*_α_* particles, denoted as *|*LD1*|*res (see Figure S2). In Figure 9, we presented the protein structure where each residue colored blue to red based on low to high value of *|*LD1*|*res. Among the residues in domains 1B, 1A, and 2A, L405 exhibits the highest discriminating value for LD1. Subsequently, we conducted an analysis of the protein sidechains around L405 for each system, revealing that residues Q404, D534, Q537, N559, and R560 show high prevalence (data not shown).

**Figure 9:**
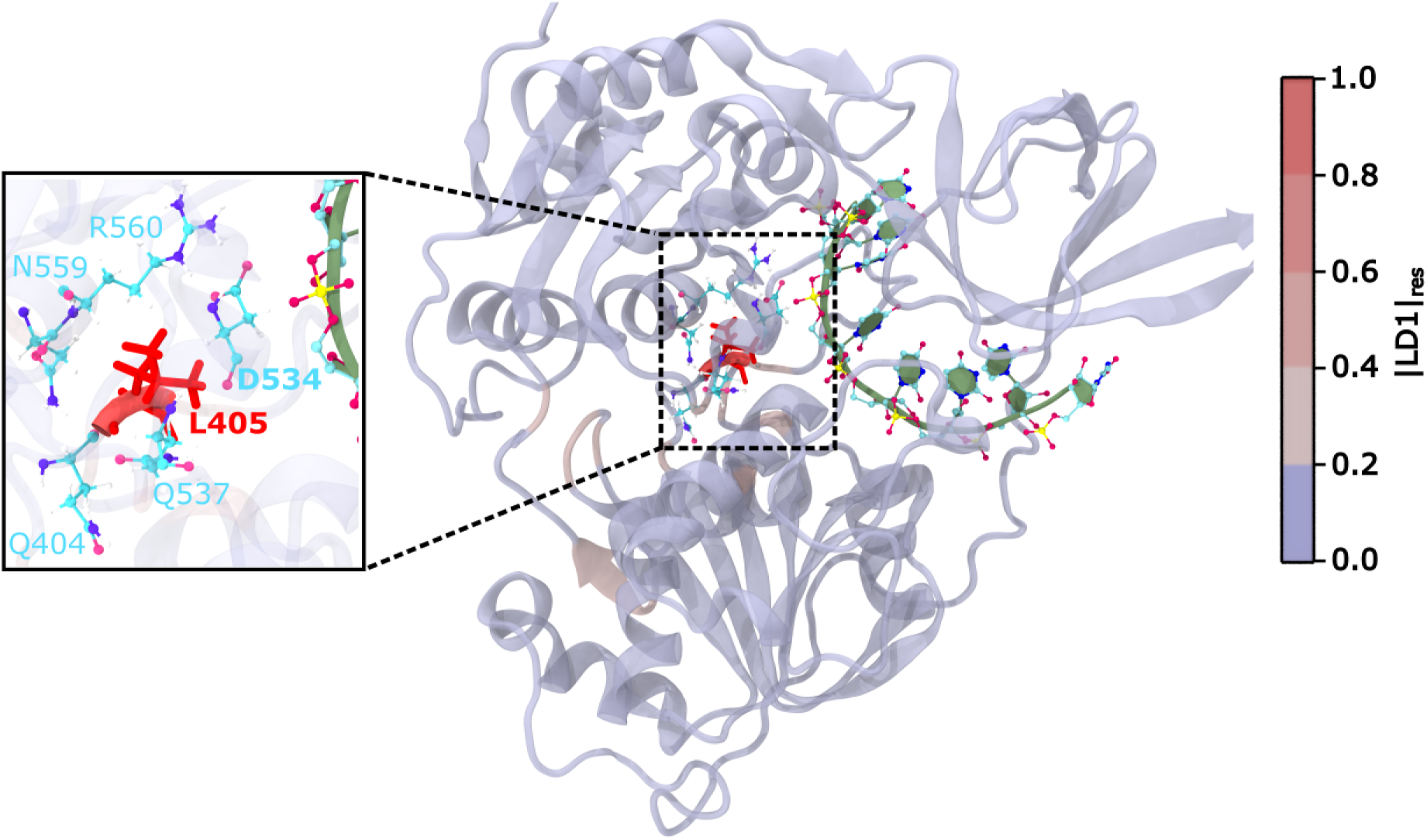
Identification of potential mutation site. In this figure, protein structure is colored according to *|*LD1*|*_res_. The LD1 denoted first discriminating vector of protein clusters and its dimension is similar to C*_α_* position. Highest magnitude (colored in red) appeared in L405. In the inset, L405 surrounding residues are shown.

Of particular interest is the behavior of D534, a motif **V** residue among the closest residues, which exhibits motion reminiscent of a power stroke. Meanwhile, Q404 belongs to motif **III** and is situated near the ATP-pocket. Our hypothesis suggests that L405, positioned at the junction between the ATP-pocket and the RNA-cleft, facilitating the transmission of the ATP-dependent effect from the ATP-pocket to the RNA-cleft. We propose that a mutation at position 405, maintaining a structurally similar yet notably altered nature, such as L405D, could generate a repulsive effect with D534, potentially competing with the D534-R460 salt-bridge, and consequently impacting the power-stroke motion.

## Conclusions

Comparative understanding of conformational ensemble of hydrolysis states delineate the underlined mechanism of translocation. In this work, we investigated the conformational heterogeneity of SARS-CoV-2 nsp13 in presence of hydrolysis substrate and products and its implications for translocation on the ssRNA tract. Our study focused on depicting the underlying processes, namely, ‘selection’ and ‘induction’, which cause alterations in the nsp13 conformational ensemble under the framework of thermodynamic cycle. We modeled ssRNA, ssRNA+ATP, ssRNA+ADP+P_i_, and ssRNA+ADP-bound nsp13 systems and conducted atomistic simulations. To accelerate the sampling of phase-space of each system, we employed GaMD simulations and leveraged the ShapeGMM clustering algorithm to identify unique conformations based on particle positions. Comparative analysis of previous modeling from our group (homology modeling based on PDB: 6JYT^21,22^) and current modeling (PDB: 7NNO^9^) qualitatively agrees with the underlying ‘selection’ mechanism at the RNA-cleft in the pre-hydrolysis states but exhibits limitations in explaining the translocation mechanism. Here, we emphasized overcoming such constraints and broadening our understanding of ATP-dependent nsp13 conformational variation by analyzing protein and RNA-cleft clusters.

Our clustering analyses on the ensemble of protein conformations comprising 1B, 1A, and 2A domains indicate four unique protein clusters. The sampling of protein clusters is synchronized with the direction of the hydrolysis cycle, manifesting a probable mechanism of hydrolysis substrate binding and product release. Interestingly, we observed population redistribution of protein clusters and the emergence of hydrolysis-state dependent clusters. Our insights suggest that the sampling of the conformational ensemble of protein clusters is underlined by both ‘selection’ and ‘induction’ processes. However, delving more closely at the RNA-cleft, the three clusters comprising motion of motifs **Ia**, **IV** and **V** relative to RNA-phosphates describes conformational ‘selection’ process as evidenced by ATP-dependent population redistribution of clusters.

Protein and RNA-cleft clusters demonstrate inchworm mode of translocation. Characterization of ^P^Cs helps us understand the transition from ‘close’ to ‘open’ (or vice versa) of inter-domain clefts mediated by the ‘mid-open’ conformation. The motion of domains display a forward movement towards the 3*′* end of ssRNA, and ^P^C2 to ^P^C4 clusters represent a downward motion of each (1A, 2A, and 1B) domain, indicating a small step translocation progression. Three RNA-cleft clusters depict loose and strong binding of motifs to ssRNA while traversing one phosphate towards the 3*^′^* end. However, a comparative view of protein and RNA cleft clusters depicts good overlap, and each protein cluster comprises at least two or more RNA cleft states. We hypothesize that while the RNA-cleft clusters fluctuate in forming loose and strong grip with ssRNA, the stepwise forward motion of domains in each protein cluster redistributes population of RNA-cleft clusters. At the end of a hydrolysis cycle, nsp13 translocates one phosphate in an inchworm stepping mechanism.

Our LD1 eigenvector of protein clusters highlights that motion of L405 plays critical role in discriminating protein clusters and suggest an important site of mutation. We propose L405D mutation will disrupt the communication of ATP-pocket and RNA-cleft followed by translocation. Our ongoing study investigates the mutation effect in hydrolysis states. Protein and RNA-cleft clusters in presence of ADP shows the emergence of ^P^C4. We speculate this observation is triggered by the release of P_i_. Inspection of the P_i_ release mechanism can help us understand this unique change, which will be the focus of our future study. We also suspect that the very low probability of ^R^C1 may be due to a sampling issue or a transient state, which can be evaluated by studying ADP release. Our study provided interesting mechanistic insights from the perspective of the conformational ensemble but excluded the motion of ZBD, stalk domains, and sidechains. It would be worthwhile to investigate their contributions.

## Acknowledgement

Research reported in this manuscript was supported by the National Institute for Allergic and Infectious Diseases of the National Institute of Health under award number R01AI166050. Computational resources for this project were provide by: (1) the High Performance Computing Center at Oklahoma State University supported in part through the National Science Foundation Grant OAC-1531128 and (2) Purdue Anvil under ACCESS project number BIO220160.

## Supporting Information Available

**Table S1:** Motifs Ia, IV, and V residues in contact with RNA “phosphates (P)” around 5.0 Å for SF1 RNA-bound helicases protein crystal structures and the percentage (%) of frames where the corresponding nsp13 residues are in contact with RNA (P) for each simulated substrate state.

| Motif | Upf1<br>(2XZL)<sup>27</sup> | IGHMBP2<br>(4B3G)<sup>41</sup> | Nsp13<br>(7NN0)<sup>9</sup> | ssRNA | ssRNA<br>+ATP | ssRNA<br>+ADP+P<sub>i</sub> | ssRNA<br>+ADP |
| --- | --- | --- | --- | --- | --- | --- | --- |
| Ia | Ser461 | Ser244 | Ser310 | 45.6 | 77.2 | 98.0 | 84.8 |
| Ia | Asn462 | Asn245 | His311 | 65.3 | 85.4 | 91.6 | 88.9 |
| IV | Pro731 | Pro540 | Pro514 | 33.2 | 7.6 | 7.4 | 28.0 |
| IV | Tyr732 | Tyr541 | Tyr515 | 92.3 | 100 | 99.8 | 97.1 |
| IV | Glu733 | Asn542 | Asn516 | 76.7 | 100 | 100 | 96.8 |
| V | Ser761 | Ser563 | Ser535 | 80.7 | 95.1 | 98.7 | 97.4 |
| V | Ala764 | Asp565 | Asp534 | 97.2 | 90.4 | 98.0 | 88.3 |

**Table S2:** Motifs I, II, III, V, and VI residues in contact with substrate (ATP), hydrolysis products (ADP + P_i_ and ADP) or Mg^2+^ within 5.0 Å for SF1 ATP-bound helicase protein crystal structures and the average percentage (%) residue-substrate contacts in the present nsp13 simulations.

| Motif | Upf1<br>(2GJK)<sup>45</sup> | Nsp13<br>(7NN0) | ssRNA<br>+ATP | ssRNA<br>+ADP+P<sub>i</sub> | ssRNA<br>+ADP |
| --- | --- | --- | --- | --- | --- |
| I | Gly492 | Gly282 | 33.8 | 28.0 | 2.4 |
| I | Pro493 | Pro283 | 99.9 | 98.3 | 83.8 |
| I | Pro494 | Pro284 | 100 | 100 | 99.0 |
| I | Gly495 | Gly285 | 100 | 100 | 100 |
| I | Thr496 | Thr286 | 100 | 100 | 100 |
| I | Gly497 | Gly287 | 100 | 100 | 100 |
| I | Lys498 | Lys288 | 100 | 100 | 100 |
| I | Thr499 | Ser289 | 100 | 100 | 100 |
| I | Val500 | His290 | 100 | 100 | 100 |
| II | Asp636 | Asp374 | 100 | 72.3 | 100 |
| II | Glu637 | Glu375 | 100 | 100 | 100 |
| III | Gln665 | Gln404 | 95.7 | 97.4 | 63.6 |
| V | Gly831 | Gly538 | 98.9 | 100 | 97.1 |
| V | Arg832 | Ser539 | 35.2 | 68.1 | 45.0 |
| V | Glu833 | Glu540 | 100 | 98.0 | 97.4 |
| VI | Arg865 | Arg567 | 100 | 100 | 96.6 |
| VI | Arg867 | Lys569 | 98.1 | 66.7 | 75.5 |

**Table S3:** Residues of motifs in which C*_α_* positions are used for distance calculations.

| Motif | Residue |
| --- | --- |
| Ia | HID311 |
| IV | ASN516 |
| V | ASP534 |

**Figure S1:**
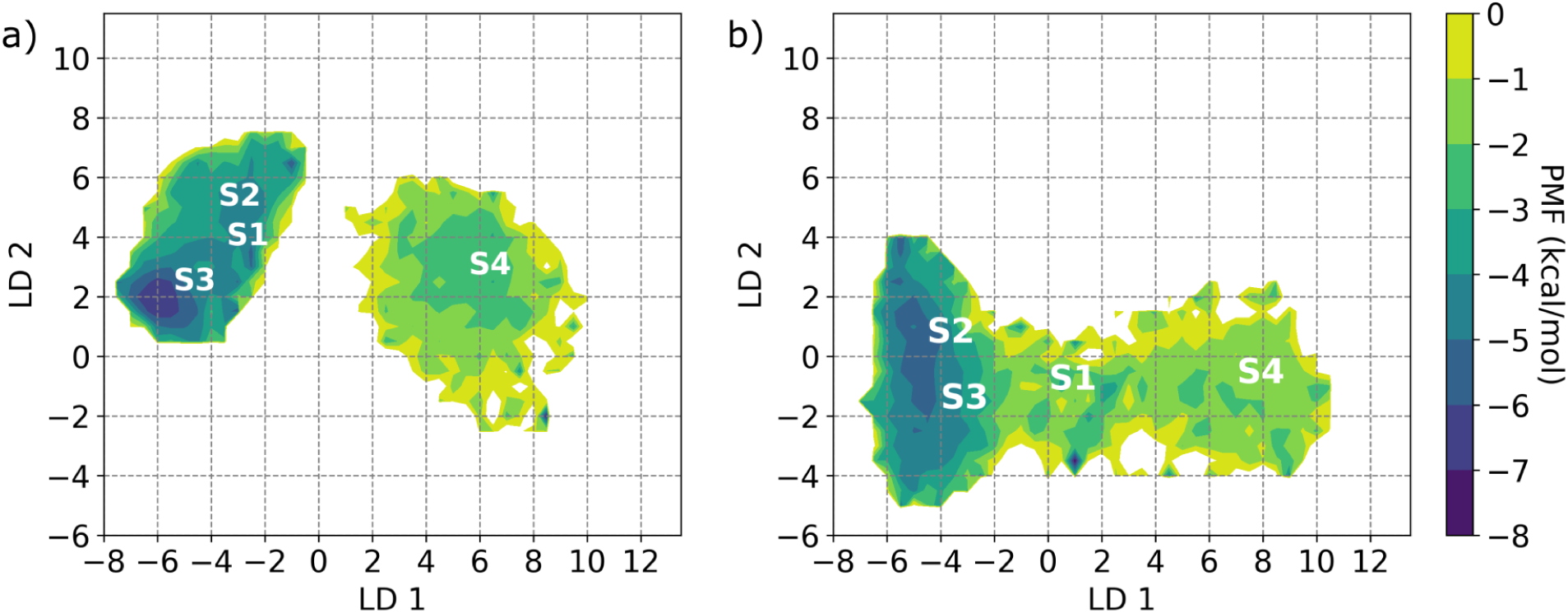
Depiction of states separability in the free energy surface. In this presentation of a)ssRNA and b)ssRNA+ATP systems, the distances of motif IV and Ia with Phosphates (P) is projected onto the LD1 and LD2 vectors. The LD vectors are generated from bGMM clustering on distance features.

**Table S4:** Average separation distance (standard deviation) between motif IV, Ia and nearest ssRNA phosphates (P) for states S1, S2, S3 and S4.

| Average Distance (Å) |  |  |  |  |
| --- | --- | --- | --- | --- |
| Residues | S1 | S2 | S3 | S4 |
| IV-P | 4.7(2) | 4.9(2) | 4.9(2) | 7.5(5) |
| IV-Ia | 19 (1) | 17.0(7) | 16.3(6) | 20(1) |
| Ia-P | 6 (1) | 7.2(7) | 5.2(3) | 9.1(9) |

**Table S5:** Probability of each states identified in GMM-LDA based clustering in ssRNA and ssRNA+ATP systems.

| Probability |  |  |  |  |
| --- | --- | --- | --- | --- |
| System | S1 | S2 | S3 | S4 |
| ssRNA | 0.04 | 0.2 | 0.5 | 0.26 |
| ssRNA+ATP | 0.04 | 0.27 | 0.5 | 0.17 |

**Figure S2:**
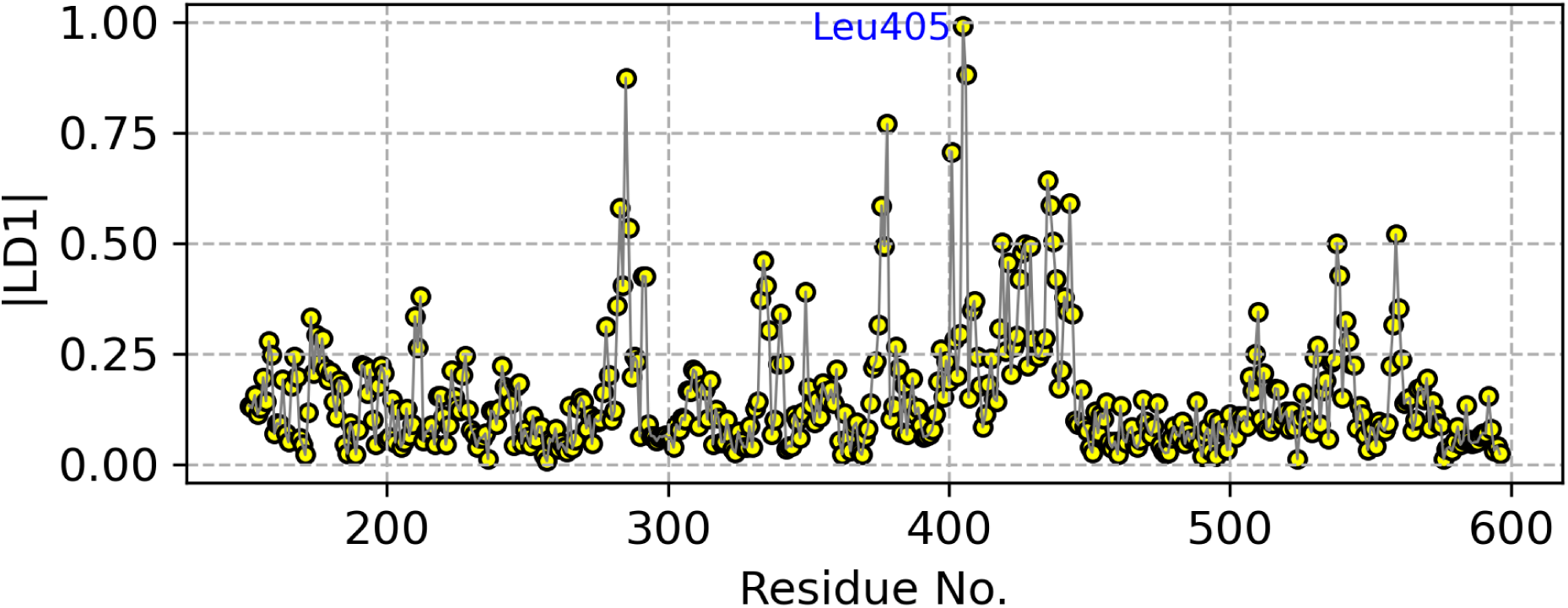
Presentation of *|*LD1*|* as a function of residue to identify important residues which best separates the clusters. In this plot the LD1 is computed from protein clusters comprising C*_α_* atoms of domain 1B, 1A and 2A.

## Notes

### Competing Interest Statement

The authors have declared no competing interest.

